# The Genomic Landscape of Prostate Cancer Brain Metastases

**DOI:** 10.1101/2020.05.12.092296

**Authors:** Antonio Rodriguez, John Gallon, Dilara Akhoundova, Sina Maletti, Alison Ferguson, Joanna Cyrta, Ursula Amstutz, Andrea Garofoli, Viola Paradiso, Scott A. Tomlins, Ekkehard Hewer, Vera Genitsch, Achim Fleischmann, Elisabeth J. Rushing, Rainer Grobholz, Ingeborg Fischer, Wolfram Jochum, Gieri Cathomas, Lukas Bubendorf, Holger Moch, Charlotte K.Y. Ng, Silke Gillessen Sommer, Salvatore Piscuoglio, Mark A. Rubin

## Abstract

Lethal prostate cancer commonly metastasizes to bone, lymph nodes, and visceral organs but with more effective therapies, there is an increased frequency of metastases to the brain. Little is known about the genomic drivers of prostate cancer brain metastases (PCBM). To address this, we conducted a comprehensive multi-regional, genomic, and targeted transcriptomic analysis of PCBM from 28 patients. We compared whole-exome and targeted RNA sequencing with matched primary tumors when available (n = 10) and with publicly available genomic data from non-brain prostate cancer metastases (n = 416). In addition to common alterations in *TP53*, *AR*, *RB1*, and *PTEN*, we identified highly significant enrichment of mutations in *NF1* (25% cases (6/28), *q* = 0.049, 95% CI = 2.38 – 26.52, OR = 8.37) and *RICTOR* (17.9% cases (5/28), *q* = 0.01, 95% CI = 6.74 – 480.15, OR = 43.7) in PCBM compared to non-brain prostate cancer metastases, suggesting possible activation of the druggable pathways RAS/RAF/MEK/ERK and PI3K/AKT/mTOR, respectively. Compared to non-brain prostate cancer metastases, PCBM were almost three times as likely to harbor DNA homologous repair (HR) alterations (42.9% cases (12/28), p =0.016, 95% CI = 1.17 – 6.64, OR = 2.8). When considering the combination of somatic mutations, copy number alteration, and Large-scale State Transitions, 64.3% of patients (18/28) were affected. HR alterations may be critical drivers of brain metastasis that potentially provide cancer cells a survival advantage during re-establishment in a special microenvironment. We demonstrate that PCBM have genomic dependencies that may be exploitable through clinical interventions including PARP inhibition.

---

Rapid Autopsy programs have identified the most common sites of metastatic prostate cancer (PCa) as the bone, lymph nodes, and the liver^1–4^. In the largest survey of over 550 autopsy cases of metastatic PCa, prostate cancer brain metastases (PCBM) were identified in only 1.5% of cases^1^. In contrast, brain metastases in other cancers are more common (e.g., 16.3% in lung, 9.8% in renal cell carcinoma, 7.4% in melanoma or 5% in breast cancer^5,6^). The recent improvement in systemic therapy for PCa has led to a significantly increased patient survival, with average survival extended to about 40 months compared to 15 months with earlier therapies (reviewed in ^5^). With prolonged survival, oncologists have noted an increased occurrence of PCBM^7^.

We posited that PCBM may require distinct genetic changes that distinguish these tumors from more common PCa metastases. To address this, we compiled a novel cohort of retrospective cases of PCBM that, when possible, included primary tumor samples before systemic therapy in order to nominate driver events. For both primary and metastatic tumors, we sampled tumors from up to three distinct regions for whole-exome sequencing (WES) and a PCa-specific targeted RNA sequencing panel. We also compared metastatic PCBM samples to metastatic PCa from other anatomic sites to provide insights into genetic alterations specifically associated with metastasis to the brain. Finally, for a subset of patients, we identified putative driver genetic events resulting from clonal evolution that could be responsible for the metastatic spread. This study is the first comprehensive genomic effort to address driver PCBM mutations.

Over the past 10 years, 10 published studies (**Supplementary Table ST1**) together interrogated 1585 metastatic PCa tumor samples using a variety of next generation sequencing approaches, most commonly WES and RNA sequencing. The metastatic sites that were interrogated included lymph node, bone, lung, and liver, but only 21 PCBM (1,3%) from 19 patients. From these 10 studies, only one included a single patient where PCBM was paired with the original primary tumor. To address this knowledge gap regarding PCBM, we devised a systematic search across hospitals in Switzerland to identify PCBM and the corresponding primary untreated PCa. We identified 28 PCBM cases (27 brain/dura and one spinal cord metastases). The primary tumor was available in 10 of these cases (35,7%). All cases were archival consisting of formalin fixed paraffin embedded (FFPE) samples (**Supplementary Data SD1**). We systematically examined the primary PCa and PCBM samples for histologically distinct areas of tumor growth with the goal of identifying heterogeneous areas for genomic analysis (Cyrta et al., manuscript in preparation, see online methods). If enough tissue available, immunohistochemistry (IHC) for the *TMPRSS2-ERG* fusion as determined by ERG overexpression^8^, and staining for *PTEN* loss and *TP53* alteration were conducted further to refine heterogenous lesions. This process identified 106 tumor areas for genomic and transcriptomic analysis (39 areas from the 10 primary specimens and 67 areas from the 28 metastatic specimens). We then performed targeted RNA on these 106 tumor areas, using a protocol amenable to FFPE samples and allowing to capture PCa-specific alterations including common gene fusion events (Salami et al.^9^) and WES. (**Supplementary Figure S1.1)**.

To compare these PCBM results to other cases of metastatic PCa, we selected the Stand Up to Cancer / Prostate Cancer Foundation castration resistant prostate cancer cohort^10–12^ (referred to hereafter as the CRPC500 cohort). This cohort is composed of 444 metastatic biopsy samples from 429 patients with publicly available WES and RNA sequencing data (**see <u>cBioPortal</u>**). We excluded tumor samples for which the anatomic site was defined as prostate (12 samples) or brain (one sample) and used the remaining 431 non-brain metastatic samples from 416 patients for comparative analysis with our PCBM cohort. To compare primary tumors from our PCBM cohort with primary tumors not selected for the advent of subsequent brain metastases, we used publicly available data of The Cancer Genome Atlas (TCGA) Prostate adenocarcinoma (PRAD) cohort, which comprises 494 patients with primary prostate cancer^13^.

In addition, we first reviewed the morphology of all 28 PCBM and 10 matched primary PCa and IHC features of all primaries and 24/28 metastases (**see methods**) to identify regions of interest for the genomic investigation. Within the cohort, pure acinar adenocarcinoma histology was identified in 25/28 (89.3%) PCBM metastases and in 10/10 (100%) patient-matched primary tumors with focal neuroendocrine (NE) differentiation identified only by IHC in 1/10 (10%). The remaining metastases 3/28 (10.7%) were classified as mixed NE carcinoma-acinar adenocarcinoma^14^ either with areas of small cell carcinoma admixed with conventional adenocarcinoma as observed in patient 27 (P27) or presenting areas with overlapping morphology between conventional adenocarcinoma and NE differentiated carcinoma (P1 and P33). This is a similar distribution of morphologic phenotypes as compared to the recent studies by Abida et al. where 11% of the CRPC cases manifested NE features^10^. The majority of the primary tumors presented high-grade areas consisting of ISUP Grade Group 5 (9/10; 90%) and Grade Group 4 (1/10; 10%). Intraductal carcinoma was present in 5/10 cases (50%). We further performed IHC analysis of proteins with frequent altered expression in PCa and observed ERG positivity in PCBM metastases and primary tumors (13/24; 54.2% and 4/10; 40%, respectively), areas of aberrant p53 expression (complete loss or gain of expression in more than 50% of tumor cells) in 20/24 (83.3%) of PCBM cases and 8/10 (80%) primary tumors). PTEN was assessed separately for nuclear and cytoplasmic expression, with cytoplasmic loss in 7/9 (77.8%, one not evaluable) and 18/23 (78.3%, one not evaluable) of the primary tumors and metastases, respectively. PTEN nuclear expression was not observed in any of the cases **(Supplementary Data SD2).**

## PCBM Cohort Mutation Summary

The 106 samples (39 from primary tumors and 67 from metastases) from 28 patients used for this study were sequenced to a median depth of coverage of 243x for primary and of 210x for metastases. Somatic mutation analysis **(see methods)** identified a total number of non-synonymous mutations (SNVs and InDels) ranging from 23 – 1820 mutations per sample (median 217 somatic mutations) (**Figure 1a**). The PCBM primary samples had a median of 184 somatic mutations per sample (range 93 - 1820), while PCBM metastases samples had a median of 259 somatic mutations per sample (range 23 - 1669). There was no significant difference in the total number of somatic mutations, SNVs alone (median 171 primary, 218 metastases), deletions (median 12 primary, 12 metastases) or insertions (median 6 primary, 7 metastases) observed between primary and metastatic samples (*q* > 0.05, Wilcoxon test) **(Supplementary Figure S1.2).** Across all samples, the highest mutation rate was consistently less than 800 mutations per sample, with the exception of patient P1, whose disease was a mismatch repair-defective PCa harboring an *MLH1* somatic missense mutation (p.Arg522Trp) coupled with the copy number loss of the other allele, with mutation rates ranging from 964 to 1820 mutations/ sample (**Figure 1a**).

**Figure 1a-d.**
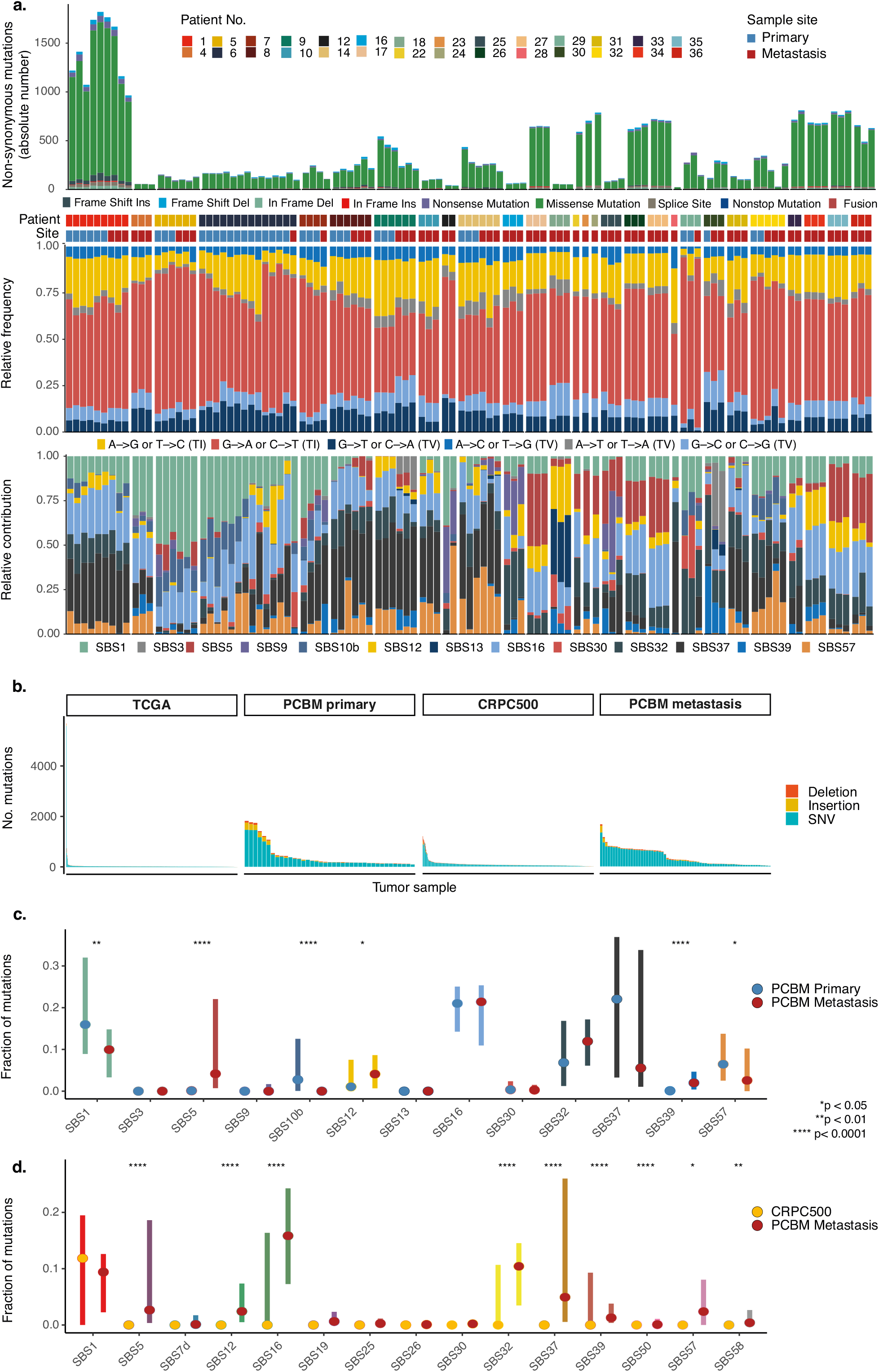
Mutational processes and number of somatic mutations in PCBM. **a** Absolute number of non-synonymous mutations (SNVs and InDels) in each PCBM sample from 28 patients (top). Relative frequency of transition and transversion mutations (synonymous and non-synonymous) (middle). Relative contribution of mutational signatures from COSMIC (bottom). **b**. Summary of total SNVs, insertions and deletions in PCBM primary, TCGA PCa primary, PCBM metastatic and CRPC500 metastatic samples. **c**. Comparison of relative contribution of mutational processes between PCBM primary and PCBM metastatic samples. **d.** As in (c) but between PCBM metastatic samples and CRPC500 non-brain metastatic samples. Points indicate median values with error bars showing the interquartile range.

Metastatic samples from the PCBM cohort had significantly higher levels of non-synonymous single nucleotide variants (SNV) (PCBM median = 218, CRPC500 median = 52, difference in mean = 263.3), insertions (PCBM median = 7, CRPC500 median = 1, difference in means = 14.8) and deletions (PCBM median = 12, CRPC500 median = 3, difference in means = 7.5) than non-PCBM metastatic PCa samples (*q* < 0.01, Wilcoxon test). Primary samples from the PCBM cohort also had significantly higher levels of SNVs (PCBM primary sample median = 171, TCGA median = 21, difference in means = 325.0), insertions (PCBM primary sample median = 6, TCGA median = 1, difference in means = 40.3) and deletions (PCBM primary sample median = 12, TCGA median = 1, difference in means = 16.5) than the TCGA primary cohort. (*q* <0.01, Wilcoxon test). The significant difference in detected mutations between primary samples from the PCBM cohort compared to TCGA primary samples, and between metastatic samples from the PCBM cohort and the CRPC500 cohort of non-brain metastases align with the high grade and stage of the tumors included in the PCBM cohort **(Figure 1b and Supplementary Figure S1.2).**

Mutational signatures were determined using a previously described approach^15^ (see methods for details). We report signatures with >15% contribution in at least one tumor sample. As expected in a population of older men (median age 71 years at time of PCBM diagnosis), the most common mutational signature was SBS1 – deamination of 5-methylcytosine, a signature associated with ageing^16^. The signature for defective homologous repair (SBS3) was observed in the metastases of two patients (P9 and P30), while it was absent in the matched primary samples from these patients^16^ **(Figure 1a)**. Substantial differences in mutational signatures were detected between primary and metastatic samples. For example, the matched primary PCa in patient P29 predominantly presented signatures for defective nucleotide excision repair (NER; SBS30), polymerase epsilon exonuclease domain mutations (SBS10b) and NER activity (SBS32) **(Figure 1c)**. In the PCBM sample from this patient, the proportionate representation of these signatures was decreased as a more complex combination of mutational signatures emerged, comprised of 16 different signatures, compared to four and nine in the primary tumor samples. There is also evidence for a change in the relative contribution of mutational processes throughout disease progression. This was best observed in patient P6, from whom samples were collected over seven years, whereby the prevalence of SBS1 (ageing) reduced throughout the period of sample collection (samples are arranged in order of collection), with a corresponding increase in the level of representation of SBS16 (transcriptional strand bias of NER activity). In this patient, SBS9 was also apparent in the latter five samples, and absent from the first six (**Figure 1a**). Mutational signatures were found to have significantly greater representation in the PCBM compared to the CRPC500 cohort (e.g. SBS5, SBS12, SBS16, SBS32, SBS37, SBS39) (*P* < 0.0001, T-test) (**Figure 1d**). Two of these signatures, SBS37 and SBS39, are of unknown etiology but have been detected in both prostate cancer and cancers of the CNS^16^. While their specific cause is unknown, SBS12, SBS16 and SBS32 have been linked to transcriptional strand bias of transcription-coupled nucleotide excision repair (NER), while SBS5 is associated with ageing, as well as mutations in the transcription-coupled NER gene *ERCC2.* These data show mutational processes distinguish PCBM from primary PCa, and PCBM from non-brain metastases, and the specific signatures showing altered representation in these comparisons implicate NER activity in this distinction.

We next explored the common molecular alterations in PCa across our study cohort in terms of gene fusions and mutational landscape (**Figure 2a, Supplementary Data SD3 and SD4**). As expected, the common truncal *TMPRSS2-ERG* gene fusion was observed in 42.8% of the 28 PCBM patients. Other common mutations, previously observed in 416 cases from the CRPC500 cohort, including *TP53*, *AR* and *RB1* were present in 28.6% (8/28) 14.3% (4/28) and 14.3% (4/28) of the PCBM cases compared to 37.9% (158/416), 13.7% (57/416) and 3.6% (15/416) in the CRPC500 cohort, respectively. There was no significant difference in the frequency of these common mutations when comparing the PCBM and CRPC500 cohorts (*q* > 0.05, Fisher’s exact test). In all cases with paired matched samples from the PCBM cohort, these mutations when present in the PCBM were always present in at least one of the primary PCa samples, suggesting the presence of early driver mutations.

**Figure 2a-c.**
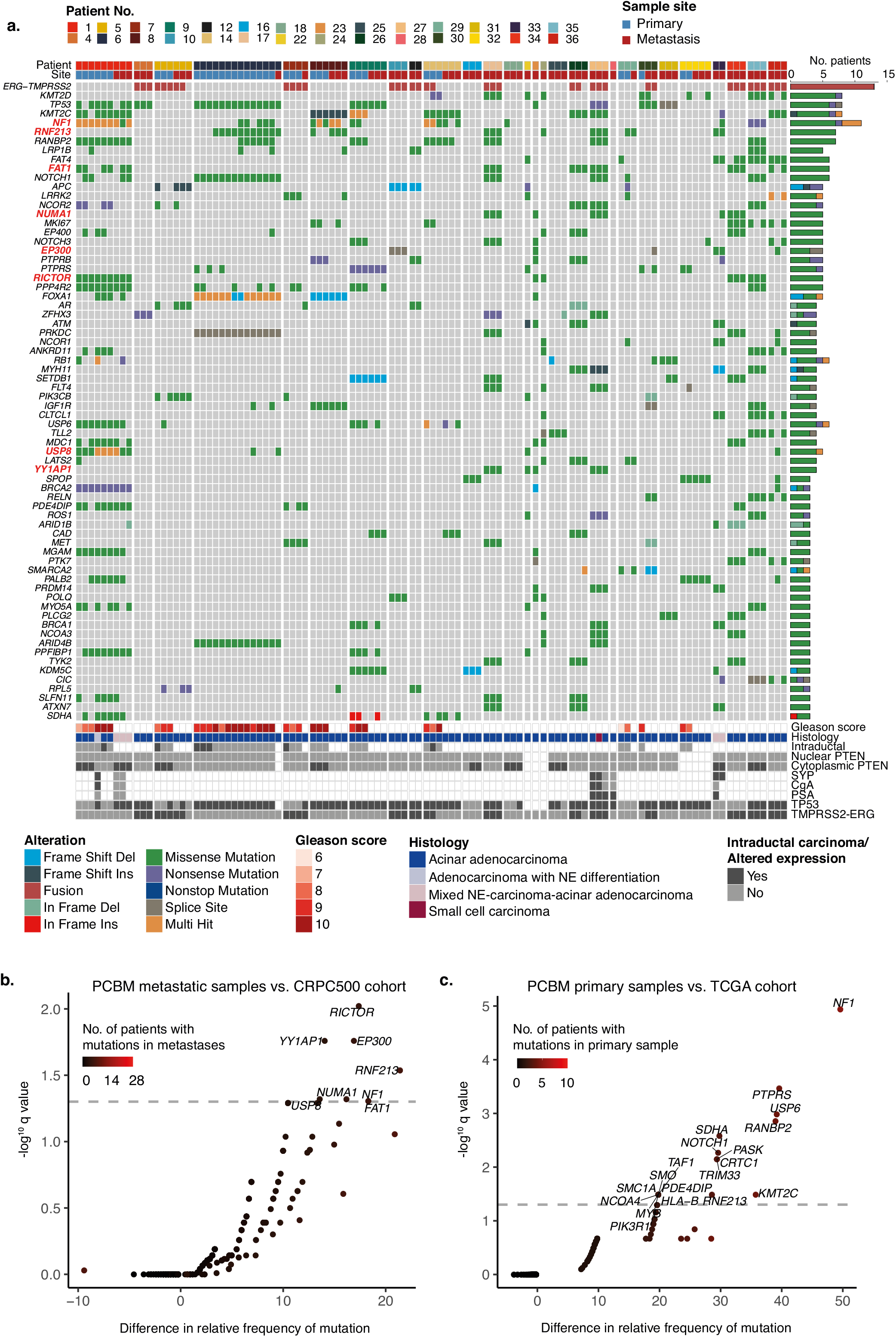
PCBM brain metastases are enriched for alterations affecting oncogenic pathways. **a.** Summary of genes showing recurrent mutation or fusion in whole exome sequencing from 106 tumor samples from 28 PCBM patients. Shown are genes with somatic alterations in at least three patients; the effects of the somatic alterations are color-coded according to the legend. Genes labelled in red are significantly more frequently altered in patients with PCBM compared to patients with non-brain metastatic prostate cancer in the CRPC500 cohort. Genes are plotted in order of mutational frequency (top). Histological assessment and immunohistochemistry results are presented (bottom). Samples are grouped by patient and sample site (primary or metastasis) as indicated above the plot. **(b-c)** Comparison of frequencies of mutations between PCBM metastatic samples and the CRPC500 cohort (**b**) and PCBM primary samples and the TCGA cohort (**c**). The x axis shows the difference in relative mutational frequency between cohorts, and the -log10(*q* value) (two-sided Fisher’s Exact Test with FDR correction) on the y axis. The color of each point indicates the number of patients harboring a mutation. Grey dashed lines indicate significance thresholds (*q* < 0.05) and genes with significant *q* values are shown.

Examining samples from patients with these mutations from whom matched primary samples were available allows assessment of the driver-status of these mutations. For example, mutations in *TP53* were found to be clonal in 10/13 primary samples from patient P6, as well as the PCBM sample from this patient. *TP53* mutations were clonal in all samples from patients P9 and P30. In patient P1 however, the *TP53* mutation was sub-clonal in both primary and PCBM samples.

Next, we asked which mutations observed in this PCBM cohort were significantly increased in frequency compared to the CRPC500 cohort, consisting of 416 patients with mCRPC in which the majority of metastatic biopsies cases from bone, lymph node, lung or liver, with only one from the brain (0.24%; this sample was excluded from comparisons)^10^ (**Figure 2b and Supplementary Table ST2**). In **Figure 2a**, we highlight in bold genes enriched for mutations in PCBM compared to the CRPC500 cohort (*q* < 0.05, Fisher’s exact test). Of particular interest was the significant enrichment for mutations observed in *NF1* (21.4% in 6/28) and *RICTOR* (17.8% in 5/28) (*q* < 0.05, Fisher’s exact test) because of their involvement in targetable pathways. These genes showed a very low mutation rate in the CRPC500 cohort with 3.1% (13/416) and 0.4% (2/416) for *NF1* and *RICTOR*, respectively. Mutations in *EP300 and YY1AP1,* genes that play a central role in chromatin remodeling, were also significantly enriched in the PCBM compared to the non-brain metastatic CRPC500 cohort with frequencies of 17.8% (5/28) and 14.3% (4/28) in the PCBM cohort and 0.9% (4/416) and 0.2% (1/416) in the CRPC500 cohort, respectively (*q* < 0.05, Fisher’s exact test) (**Figure 2b and Supplementary Table ST2**).

We also compared the primary matched tumors from the current PCBM cohort with available public data from the TCGA PRAD cohort consisting of 494 hormone naïve primary PCa tumors. Interestingly, we detected a significant enrichment for mutations in *NF1* in matched primary PCa samples from the PCBM cohort (*q* < 0.001, Fisher’s exact test), with mutational frequencies of 50% (5/10) and 0.2% (1/494) in each cohort respectively. There was no enrichment for mutations in *RICTOR* (*q* > 0.05, Fisher’s exact test), with 1/10 (10%) patients harboring a *RICTOR* mutation in the PCa PCBM samples compared to 0.4% (2/494) in the PRAD TCGA dataset (**Figure 2c, Supplementary Table ST2 and Supplementary Data SD4**).

Unlike some other common solid tumors, PCa has more somatic copy number alterations than point mutations^17^. To obtain a more complete picture of potential pathways altered in the PCBM, we interrogated the WES data for somatic copy number alterations (SCNA) in the PCBM cohort and compared our findings with the CRPC500 cohort (**Supplementary Figure S2.1 and Supplementary Data SD5**). Frequently observed SCNA in advanced PCa were detected in the PCBM cohort metastases; alterations were observed in *AR, PTEN, TP53* or *RB1*. Copy number gain or amplification of *AR* was present in 13/28 patients (46.4%). Among them 12/28 harbored those alterations in the metastases (42.8%) compared to 71.6% (298/416) in the CRPC cohort, with one patient demonstrating *AR* amplification in the matched primary tumor. One patient harbored gain and amplification in *AR* only in the primary tumor. Heterozygous loss of *PTEN* was seen in the metastatic samples of 53.6% of PCBM patients (15/28), compared to 64.7% of CRPC500 patients (269/416) and was detected in both the primary and metastatic samples in 40% (4/10) of patients for whom matched samples were available. Heterozygous loss of *TP53* was seen in the metastases of 57.1% (16/28) PCMB patients compared to 64.2% (267/416) in the CRPC500 cohort. Among patients with matched PCa primary tumor, 60% (6/10) patients showed loss of *TP53 in* both the metastasis and the matched primary samples. None of the patients demonstrated *PTEN* or *TP53* loss in their primary sample without corresponding loss in PCBM, suggesting early driver events. Heterozygous loss or homozygous deletion of *RB1* was detected in 78.6% of patients (22/28). Among them *RB1* SCNA were present in 21/28 metastases (75%), compared to 68.3% of CRPC500 patients (284/416). In six of these patients *RB1* was also found to be lost in the matched PCa, accounting for 60% (6/10) of the matched cases. In only one patient *RB1* was found lost only in the primary tumor. Eighteen PCBM patients (64.3%) harbored a concomitant loss/deletion of *RB1* and *BRCA2*. There was no significant enrichment for SCNA affecting these genes in the PCBM cohort compared to the CRPC500 cohort (*q* > 0.05, Fisher’s exact test). Interestingly *NF1*, a gene we observed to be frequently mutated, also showed frequent SCNA, with loss in 32.1% of patients (9/28). Among those *NF1* loss was detected in the metastases of seven PCMB patients (25%) although this was not significantly above the rate of 30.3% (126/416) observed in the CRPC500 cohort (*q* > 0.05, Fisher’s exact test) and in two of those patients *NF1* loss was also present in the matched primary tumors. In two other patients, *NF1* loss was detected exclusively in the primary tumor.

When compared to non-brain PCa metastases from the CRPC500 cohort, PCBM metastases showed significant enrichment for SCNA (gain or loss) affecting the PI3K pathway members *MTOR* and PI3KCD each with 67.9% (19/28) SCNA, compared to 31.2% (130/416) and 31.5% (131/416), respectively, in the CRPC cohort (*q* < 0.05, Fisher’s exact test). Of the 19 patients with SCNA affecting *MTOR* and *PI3KCD,* 5 showed CN gain; an alteration observed in 4% of cancers in a pan-cancer analysis^18^. Additionally, the tumor suppressor *LATS2,* the VEGF gene *FLT1* and the DNA mismatch repair gene *MSH3* with 71.4% (20/28), 71.4% (20/28) and 75% (21/28) SCNA, compared to 37.3% (155/416), 37.7% (157/416) and 37.5% (156/416) in the CRPC500 cohort, respectively (*q* < 0.05, Fisher’s exact test) **(See Supplementary Figure S2.2a-d and Supplementary Table ST2 for comparisons between primary and metastatic cohorts)**. *MTOR* and *PI3KCD* are both located at chr1p 36 while *LATS2* and *FLT1* are located on chr13q12, at which loci other genes were also significantly enriched for SCNA; the negative regulator of *ERBB2, ERRFI1* at chr1p36, and *FLT3, ZMYM, CDX2* at chr13q12 (*q* < 0.05, Fisher’s exact test). This demonstrates frequent copy number alterations affecting large genomic regions in PCBM. **(Supplementary Figure S2.3)**.

Taken together primary tumors and metastases from the PCBM cohort and considering mutations and SCNA, we noted 13 patients (46.2%) harboring alterations, heterozygous deletion or somatic mutations (41.9% multi hit, 48.4% missense and 9.7% non-sense) in *NF1* with five presenting CN loss without mutations, four mutation without CN loss and four presenting both. Of six patients harboring *NF1* mutations three had at least one nonsense, frameshift, or predicted deleterious mutation, as defined by MetaSVM^19^ (**Supplementary Figure S2.4**). Among the four patients with concomitant mutation and copy loss, two presented both alterations together (biallelic loss) in multiple samples from both metastases and primary tumors (P8 and P14), one in one metastasis sample (P33) and one in multiple primary tumor samples (P9). These data highlight the overall frequency of *NF1* alterations, and its potential for biallelic inactivation and, in combination with the observed mutations affecting *RICTOR* (100% missense), suggest the possibility for downstream activation of two druggable pathways, RAS/RAF/MEK1-2/ERK1-2 and PI3K/AKT/mTOR1/2, respectively.

Prior studies of CRPC have demonstrated alterations in genes involved in DNA homologous repair in up to 20% of cases^11,12,20^, making advanced PCa an important candidate for PARPi therapies^21,22^. In the PCBM cohort, 19 patients (67.8%) demonstrated alterations in *BRCA2* with heterozygous loss or homozygous deletion and in three cases with concomitant mutations. Patient P1 harbored nonsense mutations in both metastasis and PCa. Patient P23 showed a frameshift mutation and patient P36 a missense mutation within their metastatic samples. To address the status of DNA HR in the PCBM cohort, we surveyed all genes known to be involved in the HR pathway, selecting alterations used as inclusion criteria in the recently reported PROfound clinical trial^22^ (**Figure 3**). Biallelic loss (either homozygous deletion or loss of one allele together with mutation of the remaining one or 2 somatic mutations) affecting at least one of the investigated genes was found in 18% of patients (5/28). Mutations in at least one of the HR genes were detected in 39% of patients (11/28). We compared the frequency of mutation or biallelic loss affecting at least one of this set of genes detecting an enrichment for these alterations in the PCBM cohort compared to CRPC500 (*q* < 0.01, OR= 2.83; Fisher’s exact test) with a frequency of 42.8% and 20.9%, respectively. When considering the combination of somatic mutations, copy number alteration, and Large-scale State Transitions (LST), 64.3% of PCBM patients (18/28) were affected. The significant enrichment for molecular alterations affecting the HR pathway implicates PCBM patients as potential candidates for PARPi therapy in a population of men who have not been included PROfound clinical trial^22^.

**Figure. 3.**
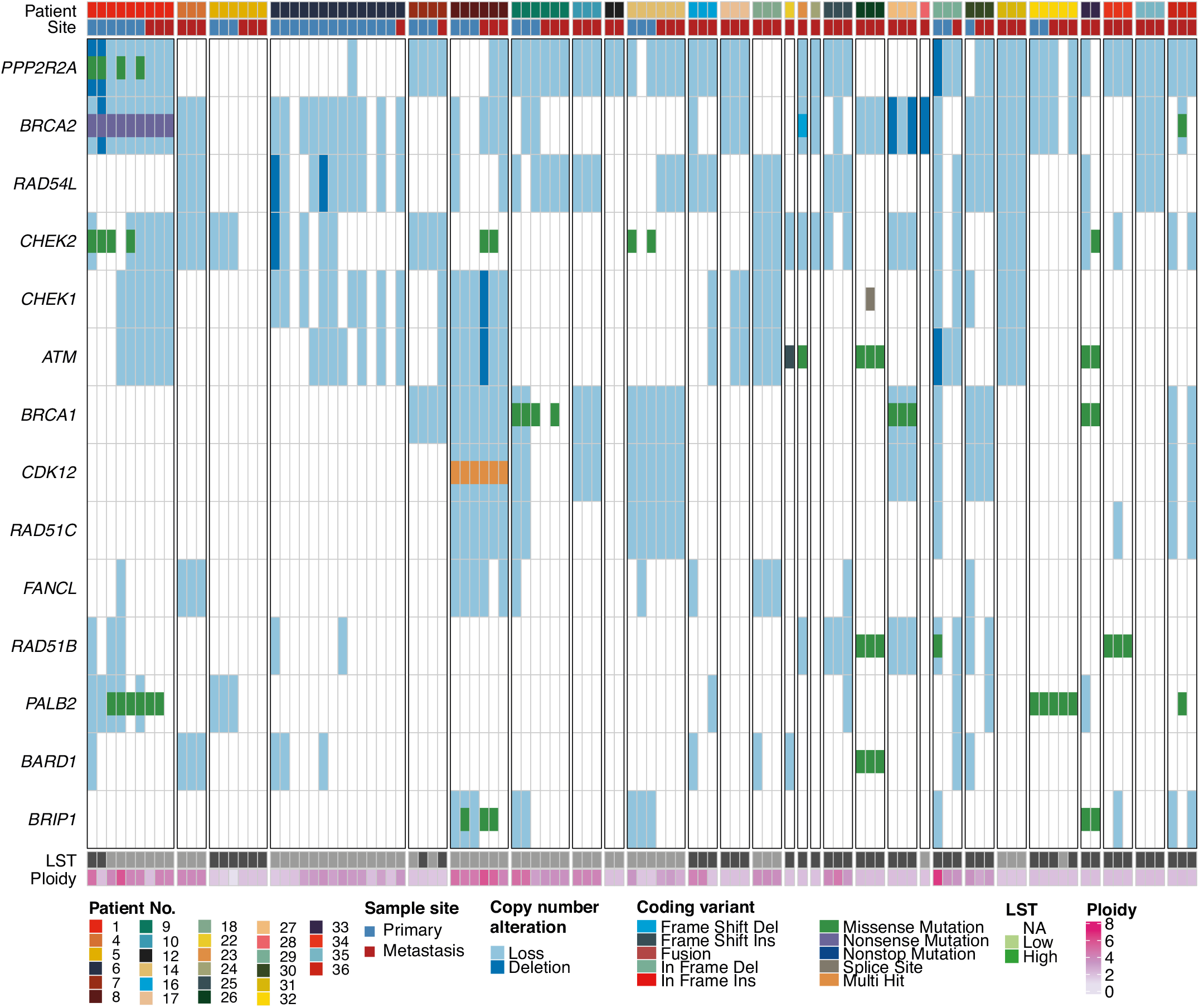
Alterations affecting HR genes are common in PCBM patients. Heatmap representing the somatic genetic alterations (mutation and copy number loss) across HR genes included in the PROfound trial, our cohort of samples (Ref 22). The effects of the somatic coding alterations (small cells) and SCNA (large cells) are color-coded according to the legend.

The multiregional approach we implemented allowed us to interrogate intratumoral heterogeneity within and between primary tumor and metastasis. First we calculated five RNA based risk scores across the cohort, including both single-transcript prognostic biomarkers (i.e. SCHLAP1 and PRCAT104) and commercially available and FDA approved prognostic scores; Myriad Prolaris Cell Cycle Progression score (CCP), the Oncotype DX Genomic Prostate Score (GP), and the GenomeDX Decipher Genomic Classifier (GC) (see Salami et al^9^). We did not find any significant correlation between prognostic scores and histological grading categories (ISUP Grade Groups) in primary tumors (Spearman’s rank p > 0.05). We also found no significant difference in prognostic scores between primary tumors and metastatic samples (Spearman’s rank p > 0.05) **(Supplemental Figure S4).**

Next, we used the cancer cell fraction (CCF) estimates for synonymous and non-synonymous SNVs from ABSOLUTE^23^ and PhylogicNDT^24^ to examine clonal evolution within and between samples from the primary and metastatic sites of patients P9 and P14 (**Figure 4**). In patient P9, following the emergence of truncal mutations including *TP53*, *SETDB1*, *PTPRS* and *KDM5C*, all three primary samples showed similar clonal architecture, with a clonal cluster of mutations (cluster 3, orange) including mutations in *NOTCH1*, *KMT2C*, *and ICK*. In the metastatic samples from this patient the prevalence of cells with cluster 4 mutations, including in *MAP3K13* (dark grey), expanded significantly from the primary samples in which the CCF of this cluster was 0.01, with these mutations becoming a clonal event in all three metastases. Despite these similarities, the metastases from patient P9 also showed substantial differences in their clonal architecture. In M1, clones with cluster 8 mutations (light pink), which included mutations in *DOT1L*, expanded to a clonal level (solid line on tree), and also expanded substantially in M3 to a CCF of 0.75 (although not reaching a clonal level). In M2 there was no such expansion of clones with cluster 8 mutations, while clones with mutations in cluster 9 (dark purple), including mutations in *MSH2* and *RPTOR* expanded to a clonal level.

**Figure 4a-d.**
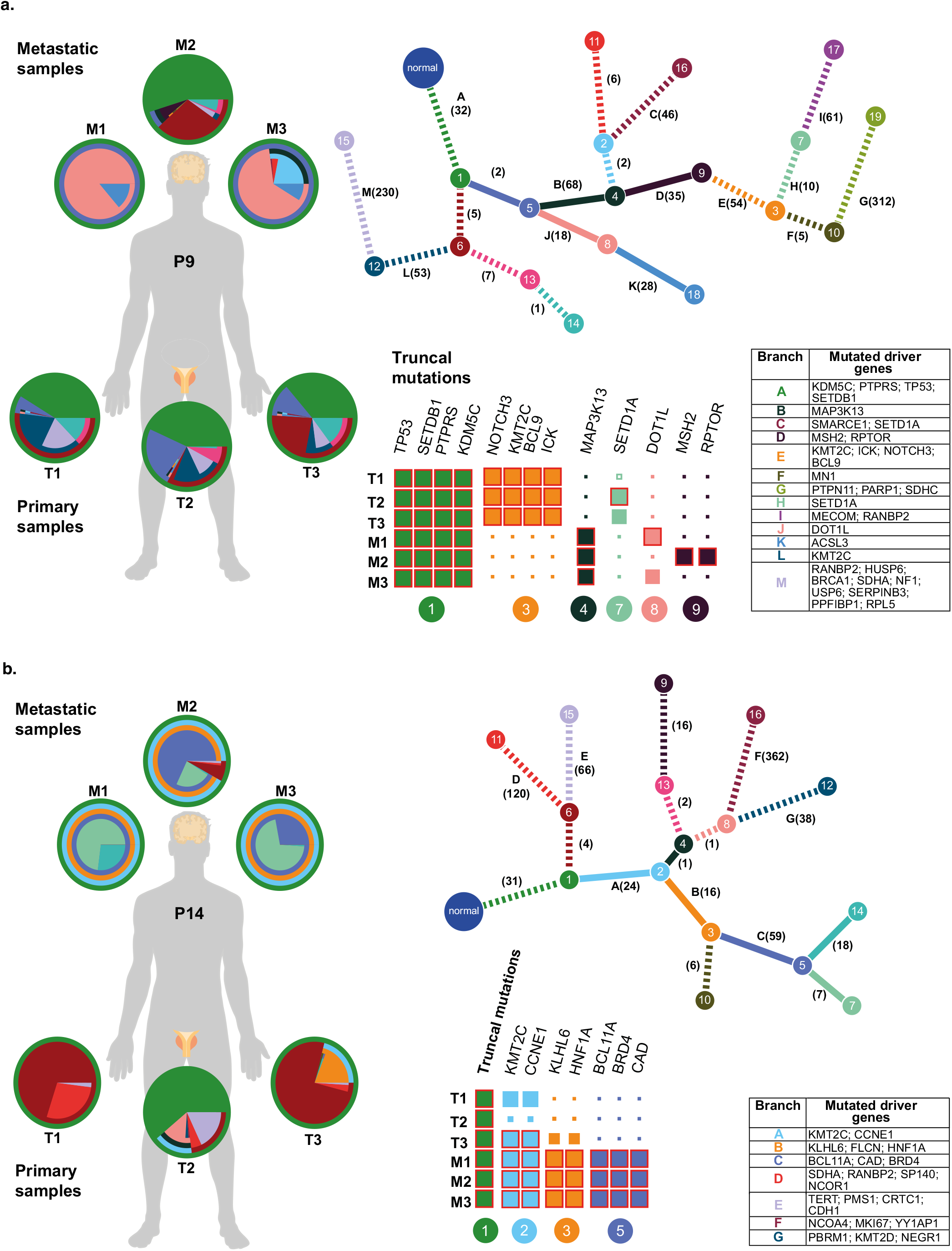
Case studies of clonal evolution in PCBM. Clonal evolution in two patients with metastatic samples from the PCBM cohort, patient P9 **(a)** and P14 **(b).** Phylogenetic trees show best solution for evolutionary relationship between clones with different clusters of mutations where each node (numbered) is a cluster of mutations. Numbers on each branch show the number of mutations distinguishing a clone from the previous (all genes) while tables show potential driver genes mutated in the distinction between a clone and the previous. Solid branches show clusters of mutations which become clonal in metastatic samples. Pie charts show the clonal composition of each sample (primary and metastatic). Segment colors correspond to branch and node colors on tree. Colored grids show CCF of mutation clusters (columns) for each sample (rows). Clusters are reduced to putative driver genes as shown in table, and the node corresponding to each cluster is labelled below. Size of squares show CCF and red outlines indicate a clonal cluster.

No mutations in putative driver genes were detected in the truncal mutations found in patient P14, however, substantial differences could be observed in the clonal evolution in this patient compared to patient P9. While the primary samples from patient P9 all showed a similar clonal architecture, primary samples from patient P14 differed significantly. In T1 and T3, cluster 2 containing mutations in *KMT2C* and *CCNE1* (light blue), was near-clonal and clonal in prevalence with CCFs of 0.84 and 0.99 respectively, while the CCF of this cluster in T2 was only 0.17, reflecting the expansion of different clones in different regions of the primary tumor. Unlike patient P9, all three metastatic samples from patient P14 showed very similar clonal architectures, with clusters 2 (*KMT2C* and *CCNE1* light blue) 3 (*KLHL6* and *HNF1A*, orange), and 5 (*BCL11A*, *BRD4*, *CAD*, purple) having a clonal prevalence in all three metastases, showing universal expansion of these clones in the metastatic setting. Clusters 3 and 5 were not present at a clonal level in any of the primary samples (cluster 3 primary CCFs: 0.01, 0.05, 0.53, cluster 5 primary CCFs: 0.01) but expanded to become clonal in all three metastatic samples (solid line in tree). In patient P14, it is possible to speculate that the likely origin of the metastases was the region from which T3 was taken, as cluster 2 (light blue) was already present to a clonal level in this sample, and cluster 3 (orange) which became clonal in the metastases had a higher CCF (0.53) than in T1 and T2 (0.01 and 0.05 respectively).

In both cases, we observed clusters of mutations, present at a subclonal level in all primary samples, becoming clonal in the metastatic samples; clones with mutation clusters 4, 8 and 9 and clones with mutation clusters 3 and 5 in patients P9 and P14, respectively. These findings are consistent with the selection for particular subclones in the metastatic niche^25^. It is, however, interesting to note that two different clones were selected for in the metastases of patient P9. In patient P9, cluster 7, while only present in a small fraction of cells in the metastatic samples (metastatic sample CCF = 0.01), expanded markedly in the primary samples to CCFs of 0.18, 0.99 and 0.78 in T1, T2 and T3, respectively. Indeed, in both patients, clones continued to emerge after the point of metastasis and clones which expanded in the metastatic setting were predicted to arise early in the phylogenetic tree (node 4 of 19 in patient P9 and node 3 of 18 in patient P14).

Taken together, these data support a model of clonal selection in the metastatic setting, and demonstrate the continued clonal evolution occurring in the primary tumor after metastasis has occurred. This implies the acquisition of mutations, which permit a cell to metastasize successfully to the brain, may be a relatively early event in the clonal evolution of these tumors.

The current study has uncovered an enrichment of genomic alterations characteristic of PCa that metastasizes to the brain. This is the first systematic approach to survey PCBM, the occurrence of which, similarly to brain metastases (BM) from other common cancers, has been increasing over recent years, due to better imaging techniques, and improvement in the overall survival of cancer patients^7^. In 2002, based on data from the Maastricht Cancer Registry, Schouten et al. reported higher occurrence of BM in patients with lung (16.3%) and renal (9.8%) cancer, compared to melanoma (7.4%), breast (5%), and colorectal cancers (1.2%)^6^.

For metastatic PCa, BM have been reported to occur in up to 2% of patients^26,27^, making PCBM considerably less common than metastases to bone, lymph nodes, lung and liver. Symptomatic BM are associated with relevant morbidity and have a significant impact on the disease course and patients’ quality of life. Current treatment options include brain directed radiotherapy, including radiosurgery, stereotactic radiotherapy and whole brain radiotherapy, as well as surgical resection. In large genomic studies over the past 10 years (**Supplemental Table ST1**), the interrogated metastatic sites have included lymph node, bone, lung, and liver, but only 21 PCBM (1.3%) from 19 patients. This low frequency may be due to the low prevalence of BM in metastatic PCa and the lack of clinical indication for performing metastatic brain biopsies. The largest prior study explored 12 PCBM using a targeted DNA sequencing approach and did not make any specific observations or comparisons between PCBM and non-brain metastatic PCa^28^. In the current study we collected and performed WES and targeted RNA sequencing on 28 cases of PCBM, the largest PCBM cohort studied to date, comparing results with a large CRPC cohort recently published by Abida et al^10^. In addition, we also identified matched primary PCa for 10 cases to explore early driver alterations and clonal evolution of tumors that eventually would metastasize to the brain. Such paired cases have been difficult to obtain due to the long-time interval between time of initial definitive therapy (i.e., radical prostatectomy) and development of metastatic PCa, and were collected from FFPE tissue. With these caveats, we asked what driver mutations emerge as specific or enriched in PCBM in the context of treated advanced PCa.

We know from several PCa Precision Oncology studies that the landscape of mutations evolves and new mutations, which are very rarely seen in primary untreated PCa, emerge during therapy^10,12,20,29–31^. For example, *AR*, *TP53*, and *RB1* mutations are enriched only following therapy. Some common mutations like the loss of *PTEN* progress from common SCNA in localized PCa to nearly universal in metastatic PCa. In the current study, we were able to address, for the first-time, which driver mutation events are enriched in PCBM compared to metastatic PCa in more common sites such as bone, lymph nodes, and liver. In a recent SU2C-PCF study exploring the CRPC500 cohort, Abida et al. failed to demonstrate significant genomic drivers determining metastatic fate to bone, lymph nodes, or visceral organs^10^. Despite the large number of patients interrogated (n=416), the CRPC500 cohort only had one PCBM. For the current study, we exploited the publicly available CRPC500 WES data as a comparator for non-brain metastatic PCa, allowing us to query for enrichment of putative PCBM driver events.

By comparing the 28 PCBM patients with 416 patients with non-brain PCa metastases from CRPC500, we discovered a significant enrichment for mutations in several genes such as *NF1*, *RICTOR*, *EP300* or *YY1AP1* compared to PCa metastases from other locations (**Figure 2b and Supplementary Table ST2)**. In particular, *NF1* alterations (copy number loss and/or mutations) were present in 10/28 (35.7%) PCMB patients. The presence of *NF1* mutations in the matched primary PCa suggests that in at least some cases this is a truncal (or early) driver event. Given the lack of *NF1* mutations in the TCGA primary PCa cohort, we propose that *NF1* truncal mutations facilitate adaptation to the CNS microenvironment as opposed to the CNS microenvironment selecting for tumors with *NF1* mutations. Future *in vivo* experiments will focus on modeling *NF1* loss and susceptibility to form metastases in the brain.

There are few studies that have to date compared brain metastases with the corresponding primary tumors. One recent study by Shih et al., explored brain metastases from lung adenocarcinoma, with the aim of nominating the specific genomic alterations associated with brain metastases^32^. Using a similar approach as the one used in our study, they performed WES on 73 cases of lung cancer metastatic to the brain and compared the results to 503 primary lung adenocarcinomas. Genomic amplifications seen in the metastatic samples compared to the primary tumors included *MYC* (12 versus 6%), *YAP1* (7 versus 0.8%) and *MMP13* (10 versus 0.6%), and significant deletions included *CDKN2A/B* (27versus 13%). They did not find an increase in *NF1* loss, but it is intriguing that *YAP1* amplifications represent another means of activating RAS signaling^33^. Shin et al. also provide *in vivo* data supporting the functional relevance of YAP1 in enabling lung cancer brain metastases. Clonality analysis performed on select cases from the PCBM cohort showed that mutations that facilitate metastasis to the brain may arise relatively early in the clonal evolution. We also see evidence that the expansion particular clones following metastasis demonstrates the selection acting on metastasizing cells in the CNS environment.

An important emerging clinical opportunity in metastatic PCa therapy is the potential use of PARP inhibition (PARPi) for common HR defects. In prior studies, the range of HR defects varied from 10 to 40% depending on the cohorts studied and definition of genes included^10–12,20,21,30^. In the current study and using a panel of HR genes associated with response to PARPi ^22^, We demonstrated HR defects are nearly three times more prevalent in PCBM than non-brain PCM. These HR alterations were also often present in the matched primary PCa samples from these patients who went on to develop PCBM, suggesting an important risk factor for disease progression. It is important to note that prior studies were unable to directly compare primaries and metastatic cases from the same patients due to the limitations mentioned above. Consistent with prior studies in metastatic PCa from various anatomic sites, the most commonly altered DNA HR genes in PCBM were *BRCA2* (5.3 – 13.3%) and *ATM* (1.6 – 7.3%)^11,12^.

As suggested by the results from the PROfound and TOPARP-B trials^22,34,35^, PARP inhibition for patients harboring *BRCA1/2* alterations, and potentially other DNA HR mutations, is an effective therapy and should soon obtain regulatory approval. The recently published PROfound phase 3 trial demonstrated prolonged overall survival (18.5 vs. 15.1 months, HR 0.64; *P* = 0.02) in the cohort of patients with mCRPC and *BRCA1*, *BRCA2* or *ATM* alterations, as well as longer radiographic progression-free survival (5.8 vs. 3.5 months, HR 0.49; 95%, *P* <0.001) in the entire cohort of included patients with mCRPC and alteration in HR repair genes, when treated with the PARP inhibitor olaparib after progression on enzalutamide or abiraterone^22^.

However, patients with known brain metastases were excluded from the PROfound trial. Testing for these alterations in advanced PCa is already being recommended by several guideline groups^36^. PARP inhibitors, such as olaparib^37,38^ or niraparib^39,40^, have shown to be active on brain metastases in patients with homologous recombination deficient (HRD) breast cancer and in murine models of brain metastases^41,42^. Moreover, HRD signatures have been reported to be enriched in breast cancer brain metastases compared with primary tumors^43^, and the occurrence of brain metastases is relatively frequent event in *BRCA1/2* mutated cancers^44,45^. In a pancreatic xenograft model, niraparib has shown higher intracranial activity compared with olaparib^41^. Rucaparib^46^ and talazoparib^47^ seem to have a more restricted blood-brain barrier penetrance. Taken together, this current study supports the likelihood that men with PCBM may benefit from PARP inhibitors given the high frequency of HR alterations.

In summary, these data suggest PCBM is molecularly distinct from advanced PCa without brain metastases, in terms of both the overall number of alterations observed, and with regard to enrichment for genetic alterations at specific genes. Although *NF1* mutations have been sporadically reported in PCa^48^, in a recent study analyzing 416 cases from the CRPC500 cohort, no *NF1* mutations were reported^10^. Our current study also identified a highly significant enrichment for alterations affection DNA homologous recombination repair genes including *BRCA2, BRCA1* and *ATM*. Both of these observations have relevant clinical implications with regards to potential therapeutic opportunities related to MEK/ERK inhibitors for *NF1* mutated patients with possible activation of the RAS/RAF/ERK signaling pathway, and PARP inhibition for patients harboring *BRCA1/2* alterations, as well as other DNA repair mutations, as recently suggested by the results from the PROFOUND and TOPARP-B trials^22,34,35^. We believe these discoveries should pave the way for future exploratory biomarker-driven clinical studies with impact on patients’ treatment.

## Methods

### Patient selection and tumor procurement

Tumor samples were collected from Pathology Departments in University and Cantonal Hospitals across Switzerland (Institute of Pathology, Bern/ Institute of Neuropathology, Zurich/ Institute of Pathology, Aarau). Inclusion criteria were defined as patients having available formalin-fixed paraffin-embedded (FFPE) blocks from confirmed CNS or meningeal metastases of prostate carcinoma and, if available, from the matched primary tumor and normal tissue (**Supplementary Data SD1**). All analyses were carried out in accordance with protocols approved by the Ethical Committee Bern (Project ID: 2019 – 00328).

### Study population

We included 32 patients across three Swiss cantons. We collected archived FFPE tissue from CNS (brain/spinal cord) and meningeal metastases with matched primary tumors in 12 cases. Most tumor samples corresponded to diagnostic biopsies (from prostate or CNS/dura), transurethral resections (TURP) or prostatectomy specimens, except for two primary tumors (patients P1 and P32) and one dura metastasis (patient P32), taken from autopsy tissue. Four patients were excluded, three due to insufficient tissue amount for WES and one due to low quantity of DNA and RNA available. For consistency and accuracy, we described the data of the 28 remaining patients, whose samples underwent both molecular analyses (i.e. WES and targeted RNA) in the manuscript. From 28/28 (100%) patients was included one metastasis and additionally 10/28 (35.8%) patients had available primary tumor tissue, with 2/10 patients including primary tumor samples at different time-points (patient P1 from two and P6 from five different time-points, respectively). In total we selected 106 tumor areas, 39 from primary tumors and 67 from metastases. IHC was conducted on 102/106 of the total areas (96.2%), including 39/39 (100%) primary tumor and 63/67 (94%) metastasis areas. All 106 selected areas underwent both molecular analyses (i.e. WES and targeted RNA) and data were obtained successfully in 106/106 (100%) for WES and in 79/106 (74.5%; 48/67 metastasis (72%) and 31/39 (79%) primary tumor areas) for targeted RNA analyses. The remaining 27/106 (25.5%) areas failed RNA analyses due to low reads depth (**Supplementary Figure S1.1**).

### Pathology review

All tissue slides (HE and IHC) were scanned and uploaded in CaseCenter (http://ngtma.path.unibe.ch/casecenter/). Through the digital microscope application CaseViewer, the slides were reviewed and annotated for further core biopsy punching. Morphological and immunohistochemical assessment was done by ARC and supervised by a board-certified pathologist (MAR). Based on the morphology, we assessed the presence of different Gleason patterns, intraductal carcinoma, ductal histology and neuroendocrine differentiation. For each specimen, we selected representative blocks to best recapitulate the heterogeneity of the above features. IHC stains were performed on all selected blocks after first review. Cases with limited amount of tissue where assessed only morphologically. For each case, p53 (clone DO-7; Dako-Agilent) PTEN (clone 6H2.1; Cascade Bioscience) and ERG (clone EP111; Dako-Agilent) were stained. Additionally, if neuroendocrine features were present, Chromogranin-A (clone DAK-A3; Dako-Agilent), Synaptophysin (clone 27G12, BioSystems) and PSA (polyclonal; Dako-Agilent) were added, as were CK5/6 (clone D5/16B4; Merck) and p63 (clone 7JUL; BioSystems) for suspected intraductal carcinoma. Finally, by combining morphological and immunohistochemical features, we identified and selected up to three heterogeneous Regions of interest (ROIs) within primary tumors and metastases. In cases showing homogeneous morphological and immunohistochemical features throughout all examined slides, up to three tumor areas were randomly selected.

### Core biopsies for genomics and transcriptomics analyses

Within the selected ROIs, core biopsies (each 1mm diameter) were annotated, punched and separately used in order of priority for targeted RNA, WES and tissue microarray (TMA)s construction.

### DNA extraction and whole-exome sequencing

After deparaffinization, DNA was extracted from selected FFPE core biopsies (1mm diameter) of matched tumor and normal tissue using the QIAamp DNA micro kit (Qiagen). Quality and quantity were determined by real-time PCR (Agilent NGS FFPE QC Kit). 10 – 200 ng of DNA underwent library preparation and exome capture using the SureSelect^XT^ low input protocol with Human All Exon V7 (Agilent) as per manufacturer’s guidelines. Multiplexed libraries were sequenced on an Illumina NovaSeq 6000 (2×100 bp) at the Clinical Genomics Lab Inselspital Bern University Hospital **(Supplementary Data SD6)**.

### Targeted RNA extraction and sequencing

Selected FFPE core biopsies (1mm diameter) of tumor tissue were subjected to DNA and RNA extraction using the AllPrep DNA/RNA FFPE kit (Qiagen). Concentrations were determined with a Qubit 2.0 fluorometer (Life Technologies). 15 – 20 ng of RNA were reverse transcribed to cDNA (Superscript VILO, Invitrogen). cDNA and 10 ng of DNA were used for library preparation with the Ion AmpliSeq Library Kit Plus with a prostate specific custom multiplex RNA^1^ or DNA panel respectively and barcode incorporation (Ion Torrent, Thermo Fisher). Template preparation of the multiplexed libraries was performed on the Ion Chef system with subsequent sequencing on the Ion S5 XL sequencer (Ion Torrent, Thermo Fisher). For each sequenced RNA sample, a coverage analysis was performed and outputted by the IonTorrent in the form of a table, providing information on the regions present in the panel, including the number of reads mapped to each region. Commercially available prognostic scores (Myriad Prolaris Cell Cycle Progression score (CCP), the Oncotype DX Genomic Prostate Score (GP), and the GenomeDX Decipher Genomic Classifier (GC) were calculated as described in Salami et al^9^.

For gene-fusion analysis for each sample all the reads that completely cover the fusion genes specific amplicons from one end to the other end (end-to-end reads) were collected and processed in order to detect the presence of fusions. Fusion genes were called when filtering criteria were met; based on the percentage of the specific end-to-end reads, the number of breakpoint reads and the presence of possible bias toward the forward of the reverse end-to-end reads **(Supplementary Data SD6)**.

### Sequence data processing pipeline and single nucleotide variant identification

Reads obtained were aligned to the reference human genome GRCh38 using Burrows-Wheeler Aligner (BWA, v0.7.12)^49^. Local realignment, duplicate removal, and base quality adjustment were performed using the Genome Analysis Toolkit (GATK, v4.1 and Picard (http://broadinstitute.github.io/picard/). Somatic single nucleotide variants (SNVs) and small insertions and deletions (indels) were detected using Mutect2 (GATK 4.1.4.1)^50^. We filtered out SNVs and indels outside of the target regions (i.e. exons), those with a variant allelic fraction (VAF) of <1% and/or those supported by <3 reads. We excluded variants for which the tumor VAF was <5 times that of the paired non-tumor VAF, as well as those found at >5% global minor allele frequency of dbSNP (build 137). We further excluded variants identified in at least two of a panel of 123 non-tumor samples, including the non-tumor samples included in the current study, captured and sequenced using the same protocols using the artifact detection mode of MuTect2 implemented in GATK. All indels were manually inspected using the Integrative Genomics Viewer^51^. To account for the presence of somatic mutations that may be present below the limit of sensitivity of somatic mutation callers, we used GATK Unified Genotyper to interrogate the positions of all unique mutations in all samples from a given patient to define the presence of additional mutations. Hotspot missense mutations were annotated using the published resources^52,53^.

### Allele-specific copy number analysis

Allele-specific copy number alterations were identified using FACETS (v0.5.6)^54^, which performs joint segmentation of the total and allelic copy ratios and infers purity, ploidy, and allele-specific copy number states. Copy number states were collapsed to the gene level using the median values to coding gene resolution based on all coding genes retrieved from the Ensembl (release GRCh38).

Genes with total copy number greater than gene-level median ploidy were considered gains; greater than ploidy + 4, amplifications; less than ploidy, losses; and total copy number of 0, homozygous deletions. Somatic mutations associated with the loss of the wild-type allele (i.e., loss of heterozygosity [LOH]) were identified as those where the lesser (minor) copy number state at the locus was 0. For chromosome X, the log ratio relative to ploidy was used to call deletions, loss, gains and amplifications. All mutations on chromosome X in male patients were considered to be associated with LOH^55^.

### SNV and SCNA enrichment analysis

Detection of genes showing enrichment for alterations over other datasets was performed using a two-sided Fisher’s exact test. Where multiple tests were carried out p values were adjusted using false discovery rate correction. An FDR (*q* value) < 0.05 was considered significant.

### Clonality of single nucleotide variants

Clonal prevalence analysis was conducted using the hierarchical Bayesian model PyClone, and the ABSOLUTE V2.0 algorithm in the case of samples used in analysis of clonal evolution. PyClone estimates the cellular prevalence of mutations in deeply sequenced samples, using allelic counts, and infers clonal structure by clustering these mutations into groups with co-varying cellular frequency. PyClone was run using a two-pass approach, whereby mutations whose cellular prevalence estimate had standard deviation >0.3 were removed before a second pass analysis was run. A cellular prevalence of >80% was used as a threshold for clonality. ABSOLUTE infers CCF from the reads supporting the reference/alternative allele, in conjunction with segmented copy-number data from WES, and was run after patching as described here: https://github.com/broadinstitute/PhylogicNDT/issues/4#issuecomment-555588341. Solutions from ABSOLUTE were manually curated to assure the solution matched the ploidy estimate generated by FACETS.

### Phylogenetic analysis

CCF histograms generated by ABSOLUTE were used as the input to PhylogicNDT^56^ to find clusters of mutations, infer subclonal populations of cells and their phylogenetic relationships and determine the order of occurrence of clonal driver events. PhylogicNDT was run using the parameters “Cluster -rb -ni 1000” to cluster and build the phylogenetic tree with 1000 iterations.

### Mutational signatures, LST and HRD

Decomposition of mutational signatures was performed using deconstructSigs based^15^ on the set of 60 mutational signatures (“signatures.exome.cosmic.v3.may2019”)^57,58^, for samples with at least 20 somatic mutations. To increase robustness, the mutations for each sample were bootstrapped 100 times and the mean weights across these 100 iterations were used. T tests were used in comparisons of the contribution of mutational signatures between datasets. Large-scale transitions (LST), a genomic marker for homologous recombination DNA repair deficiency, were computed as previously described^55^. Specifically, an LST is defined as a chromosomal breakpoint (change in copy number or allelic content) between adjacent regions each of at least 10Mb obtained after smoothing and filtering <3Mb small-scale copy number variation. Two ploidy-specific cut-offs (15 and 20 for near-diploid and near-tetraploid genomes, respectively) were used to classify tumors as “LST-high” (number of LSTs ≥ cut-off) or “LST-low” (number of LSTs < cut-off)^55^. Samples with >20% contribution by signature 3 and were LST-high were considered to display an HRD phenotype.

## Data availability

The DNA- and RNA-seq data generated through this study will be made available through a public portal.

## Acknowledgements

The authors would like to thank Mariana Ricca at the University of Bern for her expert assistance in editing and preparing the manuscript for submission.

## Authors contributions

A.R, J.G., S.G.S, S.P. and M.A.R. designed the study and the experiments. A.R and J.G. performed experiments and analysis of the results. A.R, J.G., D.A., A.F., S.G.S, S.P. and M.A.R. developed the concept. A.R. and M.A.R performed the pathology review and immunohistochemical evaluation. D.A. and S.G.S. provided a clinical perspective to the results. S.M., U.A. and V.P performed and coordinated the DNA and RNA sequencing. J.C., A.R. and M.A.R. developed the pathology review approach. A.G. and C.K.Y.N. developed modules for the bioinformatic pipeline and provided bioinformatic support. S.A.T. developed the DNA and RNA probe set design for prostate cancer. E.H., V.G., Ac.F, E.J.R., R.G., I.F., W.J, G.C., L.B., H.M. provided material and clinical data. M.A.R. provided administrative, technical and material support. A.R, J.G., D.A., A.F., S.G.S., S.P. and M.A.R wrote the initial draft of the manuscript and all authors contributed to the final version.

## Funding

Supported by an NCI (NHI) grant P50 CA211024 (S.A.T. and M.A.R.); Swiss Personalized Health Network grant SOCIBP (H.M., M.A.R.), and the Swiss Cancer League (S.P. and M.A.R.).

## Competing interests

S.A.T. and M.A.R. are co-authors (and included in the royalty streams) on patents issued to the University of Michigan and the Brigham and Women’s Hospital, on ETS gene fusions that have been licensed to Hologic/Gen-Probe Inc., who sublicensed rights to Roche/Ventana Medical Systems, and LynxDX. S.A.T. has served as a consultant for and received honoraria from Janssen, and Astellas/Medivation. S.A.T. has sponsored research agreements with Astellas/Medivation. S.A.T. is a cofounder of, prior consultant for, equity holder in, and employee of Strata Oncology. S.G.S. plays a consulting or advisory role to Astellas Pharma (Inst), Curevac (Inst), Novartis (Inst), Active Biotech (Inst), Bristol-Myers Squibb (Inst), Ferring (Inst), MaxiVax, Advanced Accelerator Applications, Roche, Janssen (Inst), Innocrin Pharma(Inst), Sanofi, Bayer (Inst), Orion Pharma GmbH, Clovis Oncology (Inst), Menarini Silicon Biosystems (Inst), MSD (Inst). S.G.S. is a co-author of the patent Method for biomarker (WO 3752009138392 A1). S.G.S. is an honorary member of Janssen and has also ties to Nektar, ProteoMediX. M.A.R. is on the Scientific Advisory Board of NeoGenomics, inc.

