## Supplementary Material for "The Genomic Landscape of Prostate Cancer Brain Metastases"

**Supplementary Figure S1.1 (related to Fig. 1)**

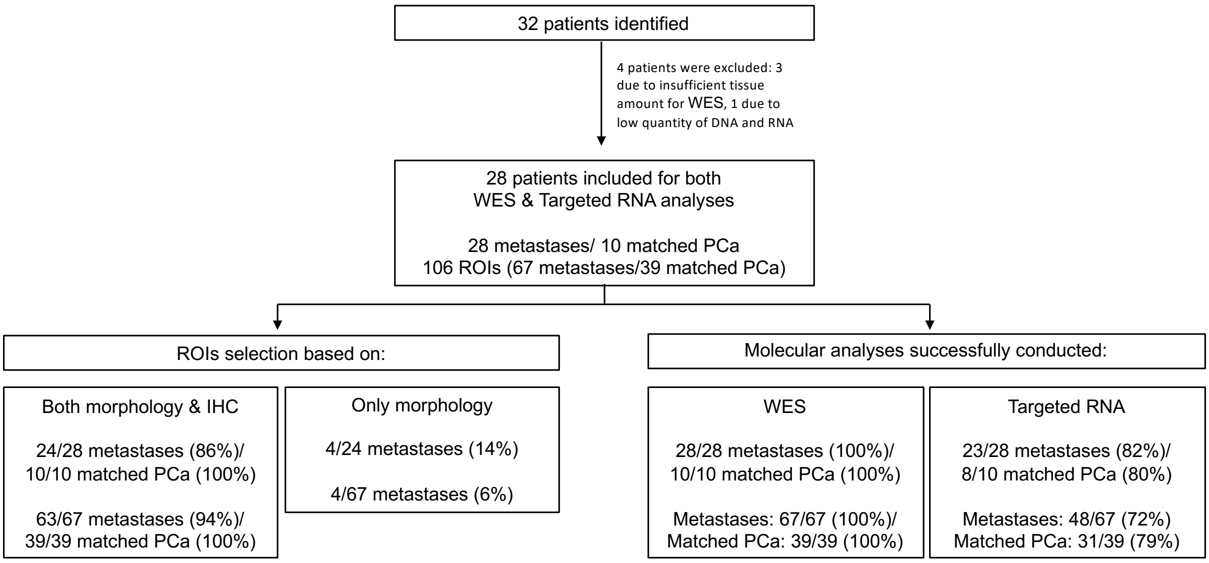


**PCBM cohort establishment and conducted methods.** Of the 32 PCa patients initially identified, four were excluded due to technical reasons. For the remaining 28 patients (described in manuscript) we selected based on morphology and, whenever enough tissue was available, immunohistochemistry the regions of interest (ROIs) within metastases and matched primary prostate cancer (PCa). All selected ROIs underwent whole exome sequencing (WES) and targeted RNA analyses. WES was successful in 100% of the ROIs. Due to low reads depth targeted RNA was only successful in 72% of the metastases ROIs and 79% of the primary tumors ROIs.

**Supplementary Figure S1.2 (related to Fig. 1)**

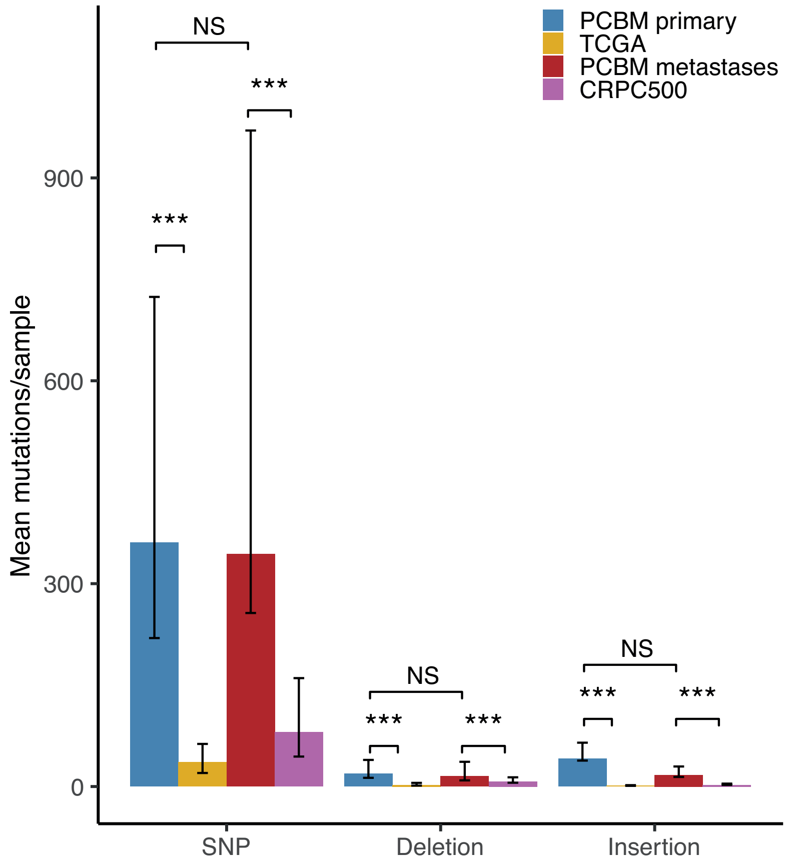


**Comparison of total mutations between the PCBM, TCGA and CRPC500 cohorts.** Bars show mean number of mutations of a given type in each cohort, and separately for primary tumors and metastases from the PCBM cohort (color coded according to legend). Values are shown for single nucleotide polymorphisms (SNP), short deletions and insertions. Error bars show the interquartile range. *** = $P < 0.001$, NS= non-significant, Wilcoxon test).

**Supplementary Figure S2.1 (related to Fig. 2)**

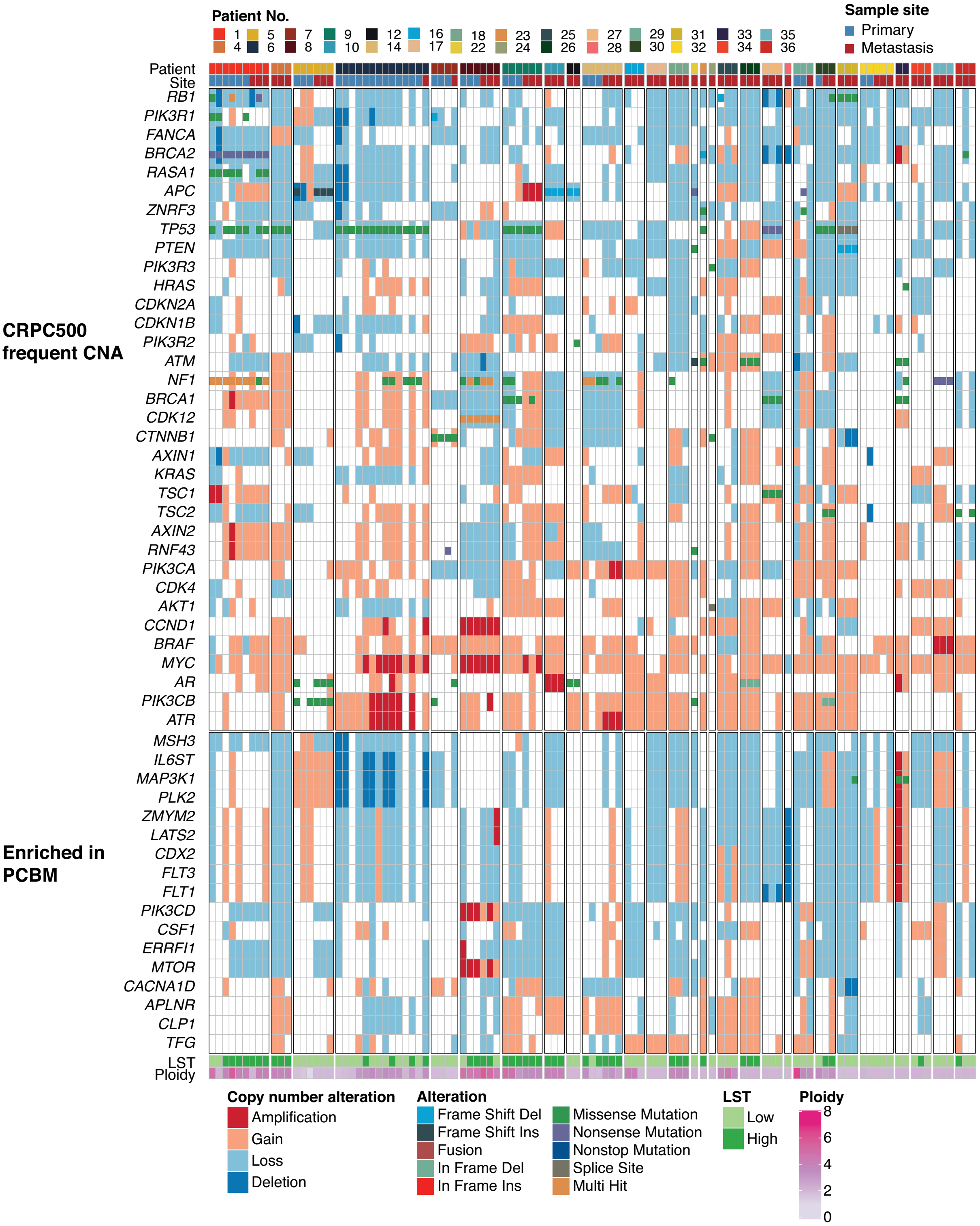


**Summary of genes with frequent and enriched SCNA.** Summary of genes showing recurrent SCNA in advanced prostate cancer (top) and enrichment of SCNA in PCBM-metastases compared to the CRPC500-cohort (bottom) from 106 tumor samples from 28 PCBM patients. SCNA are represented by the large rectangles and small squares inside rectangles represent mutations; the effects of the SCNA and coding mutations are color-coded according to the legend. Samples are grouped by patient and sample site (primary tumors or metastasis) as indicated above the plot.

### Supplementary Figure S2.2 (related to Fig. 2)

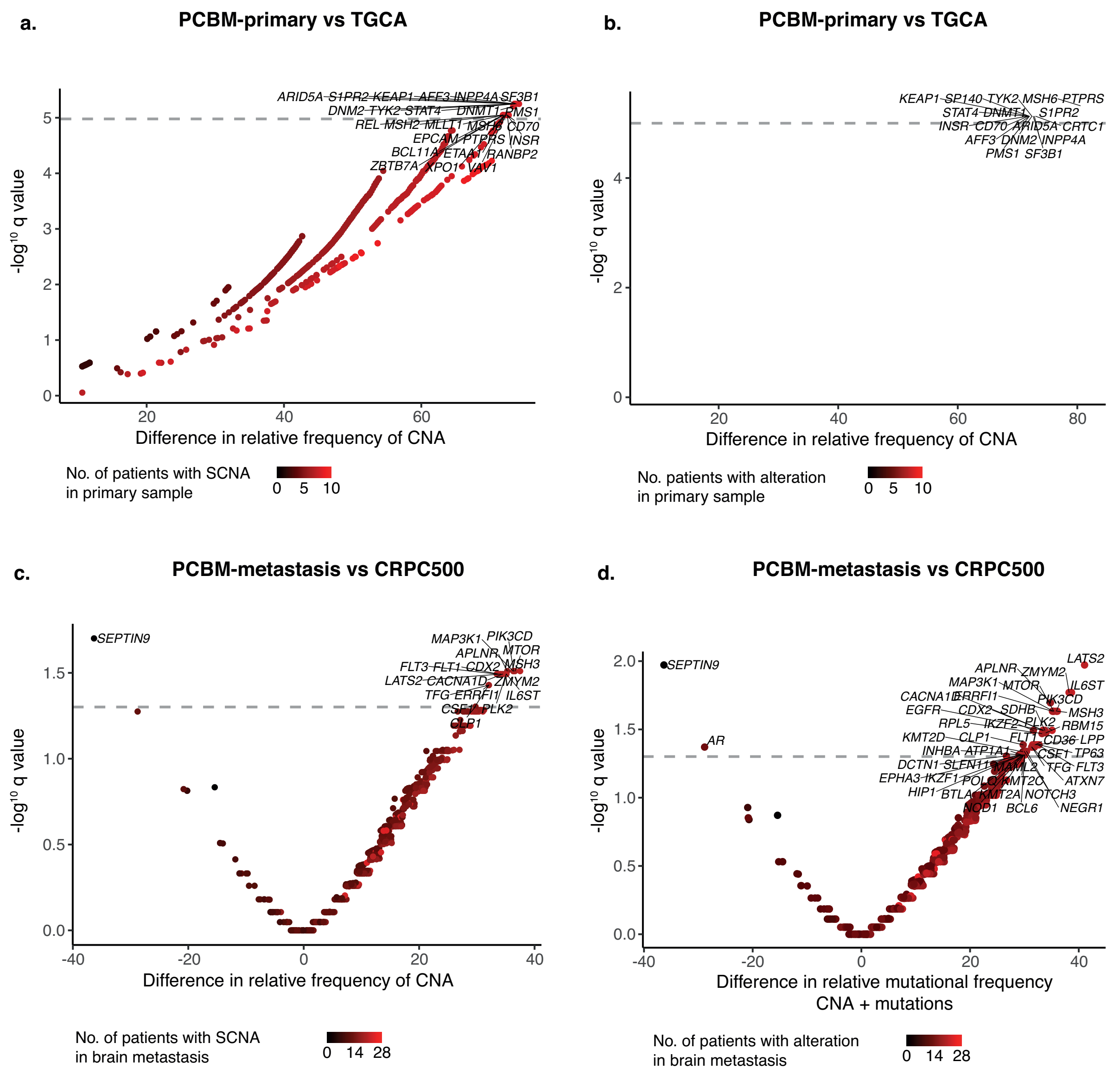


**Comparison of SCNA and any alteration (mutation or SCNA) between PCBM, TCGA and CRPC500 cohorts. (a-b)** Comparison between PCBM primary samples and the TCGA cohort of frequencies of somatic copy number alterations (SCNA) **(a)** and any alteration (mutation or SCNA) **(b). (c-d)** Comparison between PCBM metastases and the CRPC500 cohort of frequencies of somatic copy number alterations (SCNA) **(c)** and any alteration (mutation or SCNA) **(d).** The x axis shows the difference in relative mutational frequency between cohorts, and the -log10(*q* value) (two-sided Fisher's Exact Test with FDR correction) on the y axis. The color of each point indicates the number of patients harboring a mutation. Grey dashed lines indicate significance thresholds ($q < 0.05$) and genes with significant *q* values are shown.

### Supplementary Figure S2.3 (related to Fig. 2)

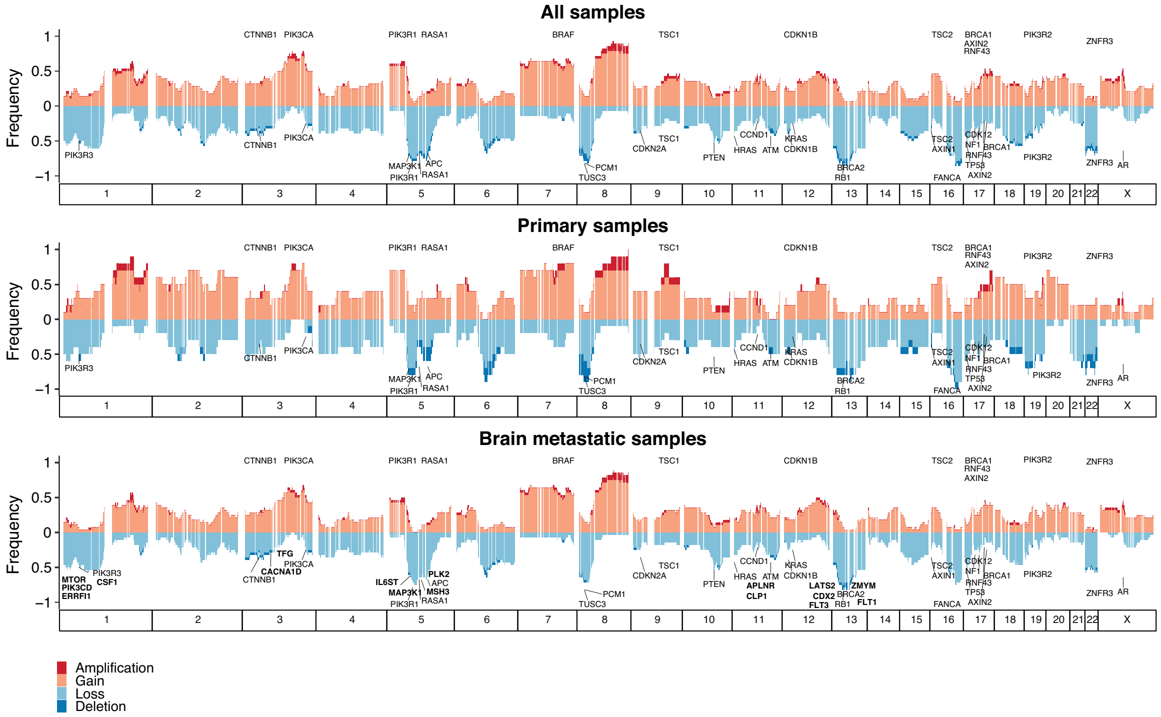


**Genome wide SCNA.** SCNA of patients (primary tumors and metastases together) (top), and separately for primary tumors (middle) and metastases (bottom). The y axis shows the frequency of gains (light red), amplifications (dark red), heterozygous loss (light blue) and homozygous deletions (dark blue). The x axis represents the locations of the alterations within the chromosomes (1-22, and X) across the genome. Frequently altered genes are labeled and genes showing enrichment for CNA in the PCBM compared to the CRPC500 cohort are labelled in bold.

**Supplementary Figure S2.4 (related to Fig. 2)**

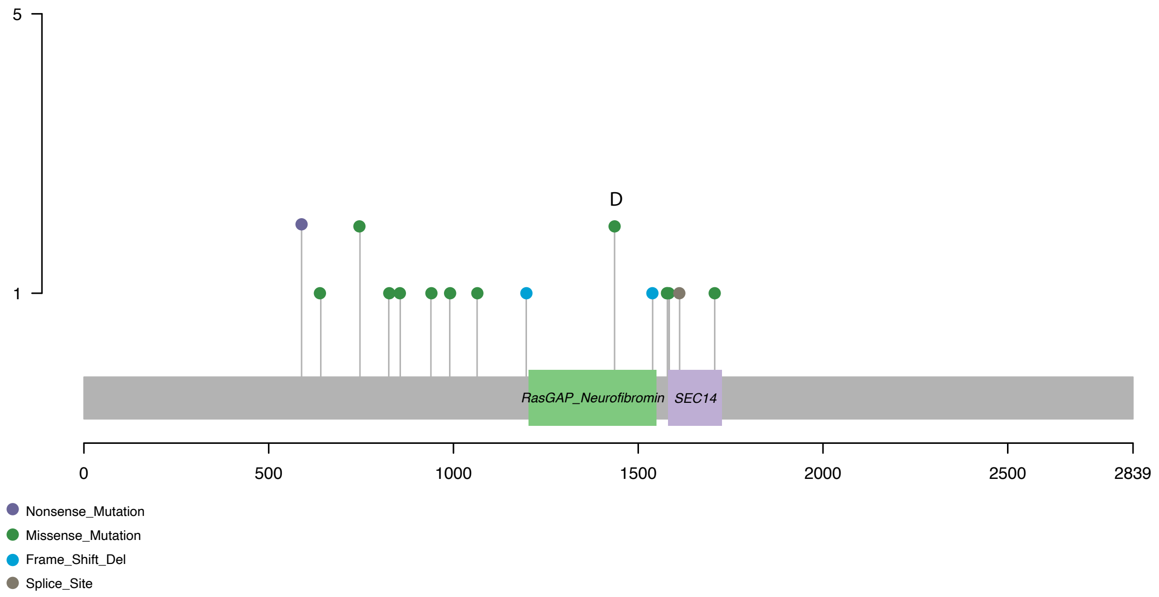


**Lollipop plot of coding mutations in *NF1*.** Height of lollipop shows number of patients with a mutation, with amino acids from the N terminal on the x axis. Mutation types are colored according to the legend. Deleterious missense mutation labelled with D.

**Supplementary Figure S4 (related to Fig. 4)**

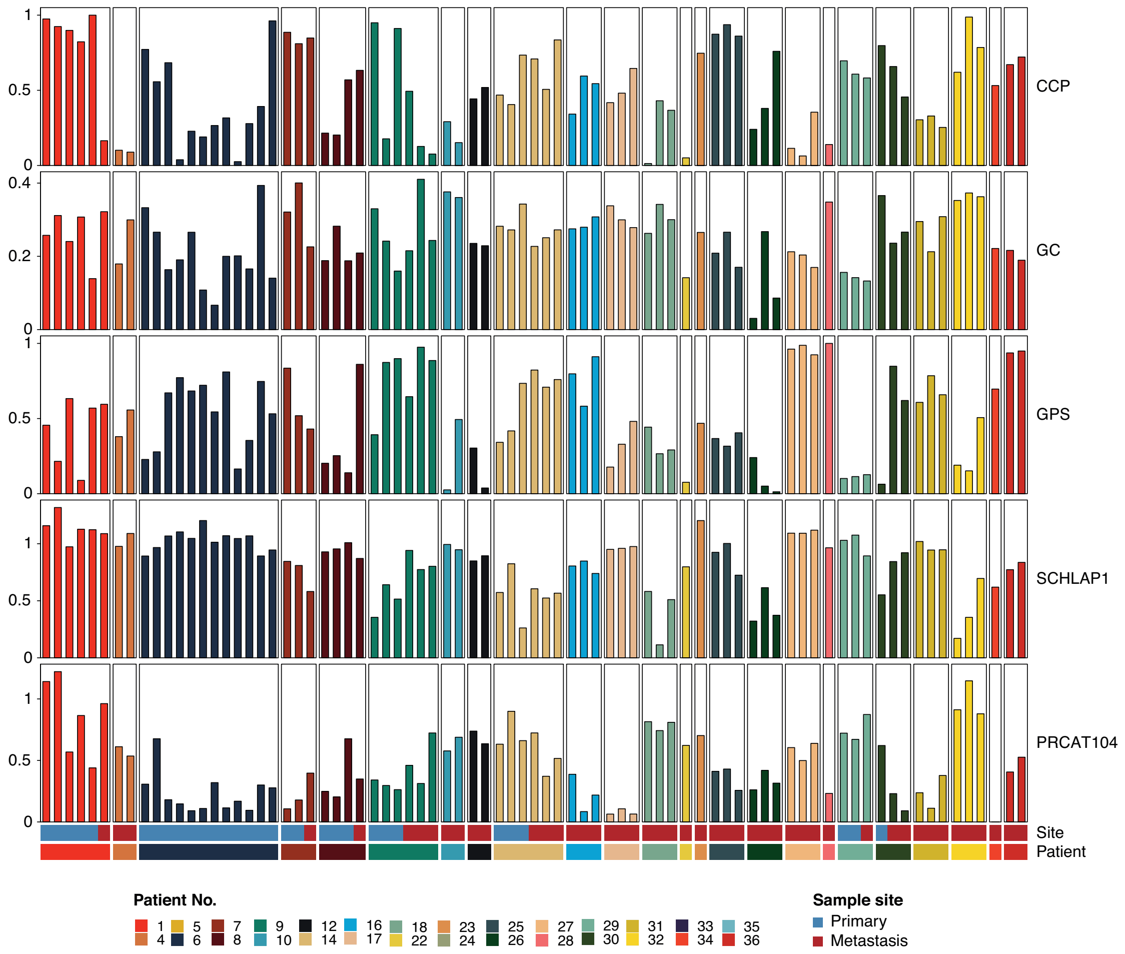


**RNA based risk assessment.** Results from five commercially available RNA based prognostic scores are shown for each sample; Myriad Prolaris Cell Cycle Progression score (CCP), the Oncotype DX Genomic Prostate Score (GP), and the GenomeDX Decipher Genomic Classifier (GC), as well as single-transcript prognostic biomarkers (i.e. SCHLAP1 and PRCAT104) (From top to bottom). Samples are grouped by patient and the source of the sample (primary or metastasis), as indicated below the plot.

**Supplementary Table ST1 (related to Fig.1)**

**Summary of large cohorts including metastases of prostate cancer within the last 10 years.** Ten large cohorts including molecular analyses of metastatic prostate cancer were reviewed based on available data from cBioPortal or publications ([1-10]. Overlapping patients were excluded (top). Patients and samples investigated in the current study (bottom). PCBM: including metastases from following locations: brain, dura (also epidural, subdural) and spinal cord. * 2 patients were previously published in Baca et al, Cell, 2013[11]. ** 150 patients were previously published in Robinson et al, Cell, 2015[12].

| | Publication | N° samples of metastatic prostate cancer | | N° patients with metastatic prostate cancer | | Methods |
|---|---|---|---|---|---|---|
| | | All locations | PCBM | PCBM | Matched primary to PCBM | |
| **1** | Taylor et al. Cancer Cell. 2010 | 37 | 12 | 12 | No | Targeted DNA/RNA-seq/miRNA expression |
| **2** | Grasso et al. Nature. 2012 | 50 | 1 | 1 | No | Targeted-DNA/RNA-seq |
| **3** | Gundem et al. Nature. 2015 | 53 | 3 | 2 | 1 | WGS |
| **4** | Beltran et al. Nature Med. 2016 | 112* | 4 | 3 | No | WES/RNA-seq/Targeted RNA |
| **5** | Kumar et al. Nature Med. 2016 | 154 | 0 | 0 | n/a | WES/RNA-seq |
| **6** | Abida et al. JCO Precis Oncol. 2017 | 228 | 0 | 0 | n/a | Targeted DNA |
| **7** | Aggarwal et al. J Clin Oncol. 2018 | 249 | 0 | 0 | n/a | RNA-seq, Targeted DNA |
| **8** | Abida et al. PNAS. 2019 | 444** | 1 | 1 | No | WES/RNA-seq |
| **9** | Van Dessel et al. Nat Commun. 2019 | 197 | 0 | 0 | n/a | WGS |
| **10** | Mateo et al. J Clin Invest. 2019 | 61 | 0 | 0 | n/a | Low pass WES/Targeted-DNA |
| | | **1585** | **21** | **19** | **1** | |

| | | | | | | |
|---|---|---|---|---|---|---|
| **11** | **Current study** | **67** | **67** | **28** | **10** | **WES/Targeted-DNA & RNA/Proteomics (GeoMx-Nanostring)** |

**Supplementary Table ST2 (related to Fig.1)**

Enriched genes in PCBM cohort when compared to CRPC500 & TCGA cohorts, separately for mutations (a) and SCNA (b) within metastases and for mutations (c) and SCNA (d) within primary tumors. Altered genes were ordered by lowest FDR. Significant mutation enrichment in the metastases was found only in seven genes (a). The Top-10 presenting the highest OR were included in (b-d).

**a**

| | MUTATIONS | | | | | |
|---|---|---|---|---|---|---|
| | PCBM-Metastases | | CRPC500 | | PCBM-Metastases vs CRPC500 | |
| Genes | Frequency (%) | N° of cases (n=28) | Frequency (%) | N° of cases (n=416) | OR | q-value |
| YY1AP1 | 14.29 | 4 | 0.24 | 1 | 66.964 | 0.017 |
| RICTOR | 17.86 | 5 | 0.48 | 2 | 43.740 | 0.010 |
| USP8 | 14.29 | 4 | 0.72 | 3 | 22.469 | 0.048 |
| EP300 | 17.86 | 5 | 0.96 | 4 | 21.914 | 0.017 |
| NUMA1 | 17.86 | 5 | 1.68 | 7 | 12.514 | 0.048 |
| RNF213 | 25.00 | 7 | 3.61 | 15 | 8.815 | 0.029 |
| FAT1 | 21.43 | 6 | 3.13 | 13 | 8.366 | 0.049 |
| NF1 | 21.43 | 6 | 3.13 | 13 | 8.366 | 0.049 |

**b**

| | SCNA | | | | | |
|---|---|---|---|---|---|---|
| | PCBM-Metastases | | CRPC500 | | PCBM-Metastases vs CRPC500 | |
| Genes | Frequency (%) | N° of cases (n=28) | Frequency (%) | N° of cases (n=416) | OR | q-value |
| MSH3 | 75.00 | 21 | 37.50 | 156 | 4.982 | 0.031 |
| APLNR | 57.14 | 16 | 21.88 | 91 | 4.740 | 0.031 |
| MAP3K1 | 75.00 | 21 | 38.70 | 161 | 4.735 | 0.031 |
| MTOR | 67.86 | 19 | 31.25 | 130 | 4.627 | 0.031 |
| PIK3CD | 67.86 | 19 | 31.49 | 131 | 4.576 | 0.031 |
| IL6ST | 75.00 | 21 | 39.90 | 166 | 4.503 | 0.032 |
| PLK2 | 75.00 | 21 | 39.90 | 166 | 4.503 | 0.032 |
| ZMYM2 | 71.43 | 20 | 36.30 | 151 | 4.372 | 0.032 |
| CACNA1D | 64.29 | 18 | 29.81 | 124 | 4.223 | 0.032 |
| LATS2 | 71.43 | 20 | 37.26 | 155 | 4.196 | 0.032 |

**c**

| | MUTATIONS | | | | | |
|---|---|---|---|---|---|---|
| | PCBM-Primaries | | TCGA | | PCBM-Primaries vs TCGA | |
| Genes | Frequency (%) | N° of cases (n=10) | Frequency (%) | N° of cases (n=494) | OR | q-value |
| NF1 | 50 | 5 | 0.40 | 2 | 220.018 | 1.15E-05 |
| SDHA | 30 | 3 | 0.20 | 1 | 191.967 | 0.003 |
| PTPRS | 40 | 4 | 0.40 | 2 | 149.818 | 0.000 |
| PDE4DIP | 20 | 2 | 0.20 | 1 | 114.675 | 0.032 |
| PIK3R1 | 20 | 2 | 0.20 | 1 | 114.675 | 0.032 |
| TAF1 | 20 | 2 | 0.20 | 1 | 114.675 | 0.032 |
| MYB | 20 | 2 | 0.20 | 1 | 114.675 | 0.032 |
| SMO | 20 | 2 | 0.20 | 1 | 114.675 | 0.032 |
| SMC1A | 20 | 2 | 0.20 | 1 | 114.675 | 0.032 |
| HLA-B | 20 | 2 | 0.20 | 1 | 114.675 | 0.032 |

**d**

| | SCNA | | | | | |
|---|---|---|---|---|---|---|
| | PCBM-Primaries | | TCGA | | PCBM-Primaries vs TCGA | |
| Genes | Frequency (%) | N° of cases (n=10) | Frequency (%) | N° of cases (n=494) | OR | q-value |
| SF3B1 | 80 | 8 | 6.54 | 31 | 55.785 | 5.40E-06 |
| INPP4A | 80 | 8 | 6.96 | 33 | 52.230 | 5.40E-06 |
| AFF3 | 80 | 8 | 7.17 | 34 | 50.608 | 5.40E-06 |
| STAT4 | 80 | 8 | 7.17 | 34 | 50.608 | 5.40E-06 |
| KEAP1 | 80 | 8 | 7.17 | 34 | 50.608 | 5.40E-06 |
| DNM2 | 80 | 8 | 7.17 | 34 | 50.608 | 5.40E-06 |
| ARID5A | 80 | 8 | 7.17 | 34 | 50.608 | 5.40E-06 |
| TYK2 | 80 | 8 | 7.17 | 34 | 50.608 | 5.40E-06 |
| S1PR2 | 80 | 8 | 7.17 | 34 | 50.608 | 5.40E-06 |
| DNMT1 | 80 | 8 | 7.38 | 35 | 49.077 | 5.95E-06 |

**Supplementary Data (see excel file in Data Sets)**

**SD1.** Patients and tissue information

**SD2.** Regions of interest (ROIs) selected including samples description, methods conducted and pathology assessment (morphology and immunohistochemistry) (n=32 patients)

**SD3.** Mutation call for PCBM-metastases and CRPC-cohort and comparison between both

**SD4.** Mutation call for PCBM-primary tumors and TCGA-cohort and comparison between both

**SD5.** SCNA call for PCBM-metastases and CRPC-cohort and comparison between both

**SD6.** Summary of the protocols used for DNA and RNA extraction, library preparation and sequencing for the different methods
